# An Artificial Intelligence-based framework for protein interaction design with accelerated KAN-based Positive-Unlabeled learning

**DOI:** 10.64898/2026.01.28.702421

**Authors:** Shubhrangshu Ghosh, Pralay Mitra

## Abstract

Protein design seeks optimal amino acid sequences for target structures, but designing stable protein complexes remains challenging. We introduce a protein interaction design pipeline combining Monte-Carlo simulation with Metropolis-criteria (MCM) and Deep-Learning. It uses Protein-Protein-Interaction(PPI) scores from Deep-Learning-based MaTPIP model to form a PPI-score-based-MCM (PMCM). The workflow integrates PMCM-driven sequence generation, HDBSCAN-clustering-based selection, and validation via AlphaFold2 and Molecular-Dynamics (MD) simulation. Incorporating learned PPI scores enhances efficiency and feature fusion. A Positive-Unlabeled (PU) learning classifier accelerates sequence validation, while the Kolmogorov-Arnold Network (KAN) improves PU learning over Multi-Layer-Perceptron (MLP). AlphaFold2 predictions yield median Root-Mean-Square-Deviation (RMSD) 1.17 *Å*, predicted-Template-Modelling (pTM) 0.72, and interface pTM 0.88; 73% of complexes remain within 2 *Å* RMSD after 100 ns MD, confirming stability. Interface mutations reveal altered interactions. The KAN-based PU model improves F1-score, precision, and AUC by 5%, 11%, and 2% over MLP. Overall, our method outperforms traditional and simulation-based methods while remaining competitive with modern Deep-Learning design frameworks.

## 1 Introduction

Protein design is a non-deterministic polynomial-time hard problem with no analytical solution. The two key components of the protein design method are a sequence-structure compatibility measure (like an energy function [23, 12, 9]) and a search strategy for guiding residue mutations in the sequence space. Stochastic search strategies, such as Monte Carlo (MC) simulations [24, 7, 20], are computationally efficient and frequently used to predict amino acid sequences that best fit a target protein structure. Recently, TIMED [4] presented a data-driven Convolutional Neural Networks (CNNs)-based protein sequence design method.

GENERALIST [1] is a generative model-based protein design approach that offers simpler learning, greater tunability, and higher accuracy compared to other generative methods. It predicts locally optimal, conservative sequences that are likely to fold into stable 3D structures. Two recent protein design approaches using MCM are Modularity-based parallel protein design [24] and SADIE [20]. The first method used protein-unit modularity and parallel MCM for the effective sequence search in protein design, while the second introduced a greedy simulated annealing MC algorithm for efficient sequence search convergence.

In this work, for the first time, we demonstrate that a Deep Learning-based PPI model and its PPI confidence score (MaTPIP-Hybrid [10]), as a stability metric, can replace the energy function in a novel form of Monte Carlo simulation with Metropolis criterion (MCM) to achieve improved accuracy at a faster speed in the protein interaction design pipeline. We have integrated all these ideas in the present work and proposed a novel usage of Kolmogorov-Arnold Network (KAN)-based Positive-Unlabeled (PU) learning. The PU learning-based approach [2, 19, 28] is applied here to expedite certain stages of the protein interaction design pipeline, with the KAN [18, 11] classifier utilized in both stages of the PU learning process: first, to identify reliable negative examples, and then to train a supervised classifier on the dataset comprising positive and reliable negative samples. This represents a unique application of KAN in PU learning for protein complex design. No prior studies have adopted an MCM approach to explore the sequence search space using the PPI score or proposed KAN-based PU learning for replacing resource-intensive, time-consuming post-simulation processing to select final designed candidates ready for the experimental work. In comparison with other methods, it is evident that our pipeline convincingly exceeded conventional evolutionary or the latest simulation-based methods and was highly comparable with the recent data-driven, Deep Learning-based protein design approaches.

## 2 Method

The architectural flow diagram of the proposed framework is shown in Fig 1. The different components shown in this figure are described below.

**Fig. 1.**
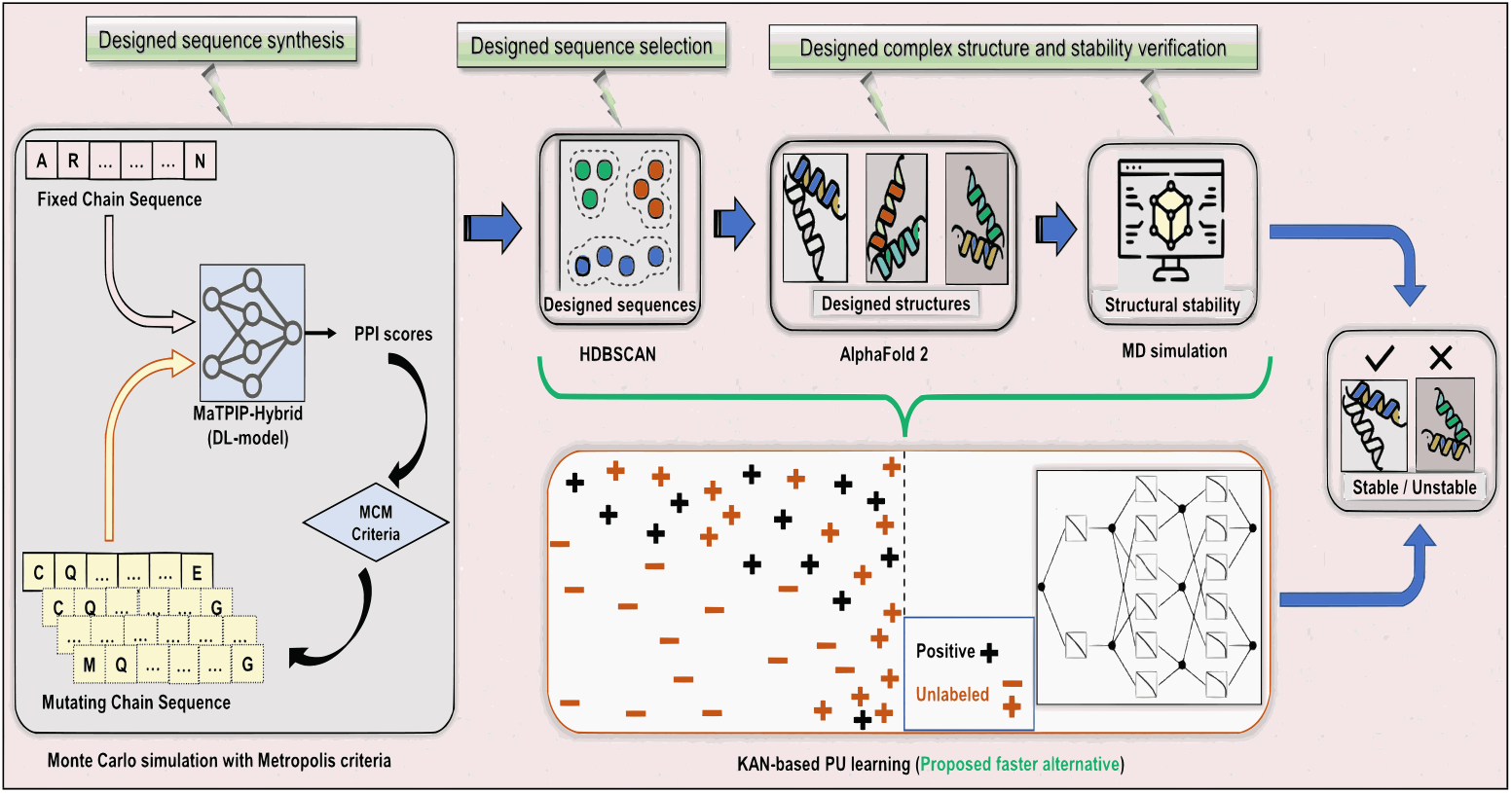
Overall architectural flow of the proposed framework

### 2.1 PPI score-based MCM (PMCM)

#### Dataset

We utilize the Protein-Protein Docking Benchmark v5.5 [26, 13], a widely used dataset with experimentally determined protein complexes and their unbound structures. We randomly select 69 representative dimers, encompassing a range of lengths (from 128 to 977 residues), and diverse fold classes as per Structural Classification Of Proteins. During design, we fixed the first chain sequence, while the other chain undergoes amino-acid mutation(s) at some randomly selected position(s). Here, we use the terms *mutated* and *designed* interchangeably.

We generate a large number of designed sequences (typically, 30000) through MCM iterations. In each iteration, at least one random mutation is performed on the previous sequence and the MaTPIP-Hybrid [10] PPI score is predicted for the designed complex. Instead of the conventional energy-based MCM approach, here, we have taken a unique approach (PMCM), where the probability is derived from the PPI confidence score predicted by MaTPIP-Hybrid. The PPI score in MCM is superior compared to the energy-based calculation since MaTPIP-Hybrid infers this PPI score based on various features, including structure-aware embeddings. The embeddings from multiple pre-trained Protein Language Models (PLMs), like ProtTrans [5] (ProtT5) and Bepler & Berger’s model [3] which is a protein structure-aware model are used. The manually curated features, including position-specific scoring matrix (PSSM), LabelEncoding, SkipGram, Blosum62-based features, and one-dimensional Autocovariance, Pseudo Amino Acid Composition, Conjoint Triad Method, and Local Descriptor are also used for this inference. Grounded in protein sequence data, the PPI score emerges as one of the most effective scoring schemes in protein engineering. Higher PPI scores indicate greater stability of the protein complex, motivating our approach.

Clearly, in MCM, the PPI score-driven approach works opposite to the energy-based method — higher PPI scores are better, unlike lower energy values. Thus, we redefined the acceptance criteria of the PMCM. The original chain-pair of the native complex has a PPI score *>* 0.5, and our goal was to make this PPI score to roam around at the higher level through the iterations (involving mutations) of MCM. We also allowed the occasional perturbations of this trend using the Metropolis criterion for the better exploration of the sequence search space and to avoid any local optima. Throughout the PMCM process, the number of mutations per iteration is at most, 10 or 10% of the current sequence length, whichever is lesser. Mutated sequences in PMCM were accepted if their PPI score is at least 0.5.

##### Algorithm 1: Protein Complex Design

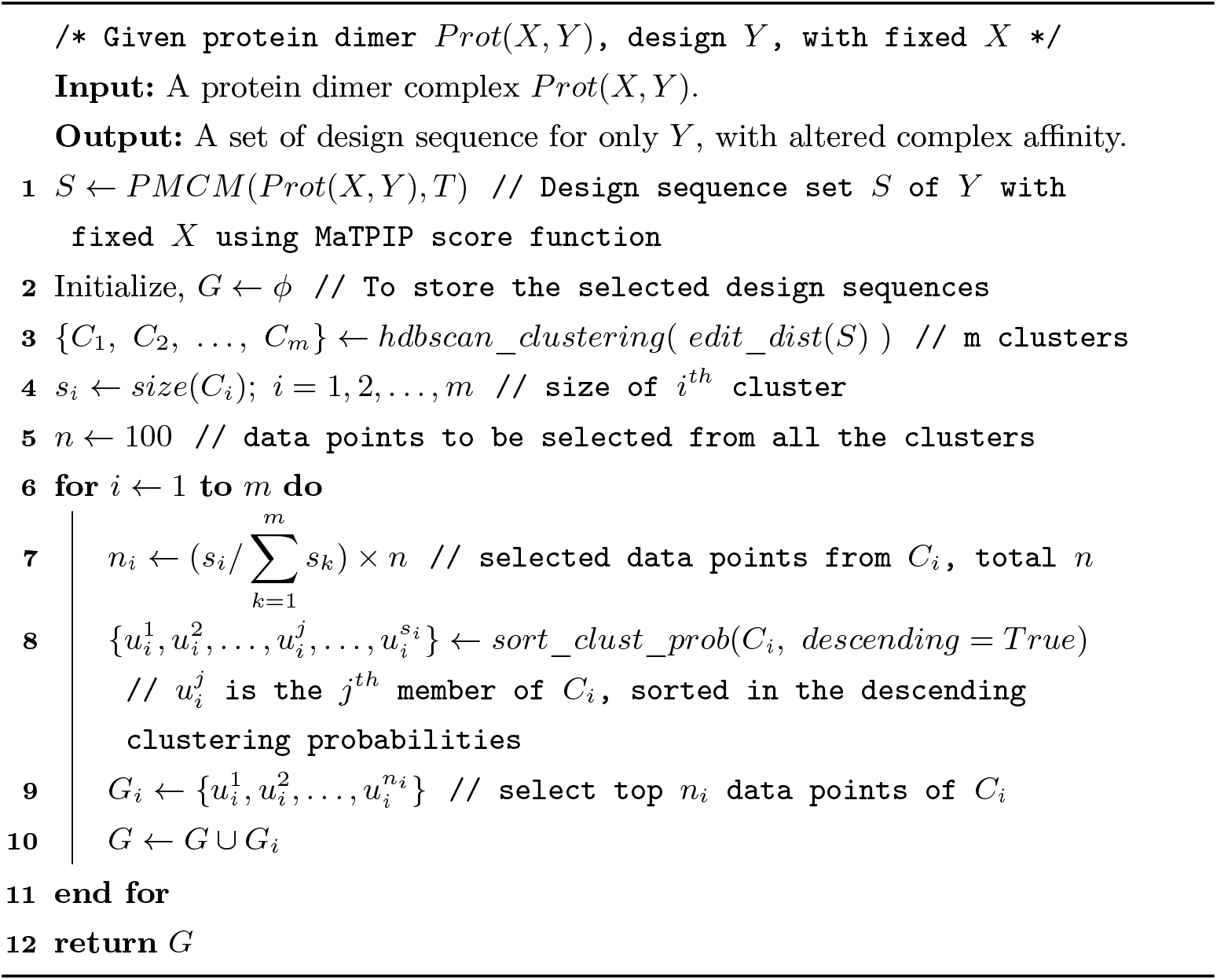

#### Edit-distance based HDBSCAN clustering

The PMCM resulted in a large number of designed chain sequences (accepted during simulations) for each chain-pair of each protein complex. To analyze these for the final design sequences, we applied edit-distance-based HDBSCAN [25, 21] clustering. We calculated the edit distance (negative of the BLOSUM62-based similarity score) between each sequence pair to create a distance matrix for clustering. HDBSCAN automatically determined the number of clusters and identified outliers, which were removed. We then selected a set of 100 sequences from all clusters proportionally to their sizes based on descending clustering probabilities. For example, if three clusters had sizes of 3000, 2000, and 5000, for a specific chain-pair of a protein complex, we would select 100 data points in the ratio of 3:2:5, resulting in 30 from cluster 1, 20 from cluster 2, and 50 from cluster 3, each chosen based on clustering probability. These selected sequences were then subjected to PSI-BLAST, keeping only those with *<* 70% similarity to avoid distant homologs. The complete procedure for generating and selecting the designed sequences is outlined in Algorithm 1. Our analysis reveals that a significant number of interface residues were also mutated, suggesting alterations in the interaction (see *Interface residue mutation for altered interaction* later).

### 2.2 Verification of the selected sequences

#### AlphaFold2-based structure prediction

The selected designed chain sequences from the previous step were input into AlphaFold2 [14] for structure prediction, with side chain relaxation using AMBER. Each predicted structure was compared with its native counterpart and only sequences with an RMSD below 12 *Å* were retained. Next, the qualified designed chains were concatenated with the fixed member of the corresponding chain-pair to form the designed complexes, which were then input into AlphaFold2’s multimer model [8] for complex structure prediction. The RMSD between designed and native complex structures was calculated, with a threshold set at 8 *Å*. The eligible designed complexes were sorted by ascending RMSD. We utilized a local implementation of ColabFold [22] referred to as LocalColabFold. AlphaFold2 is computationally intensive, while more lightweight and faster alternatives like OmegaFold [27] and ESMFold [15] exist, we chose to use AlphaFold2 due to its superior performance [6] in protein structure prediction.

#### MD simulation-based structural stability verification

The structural stability of the designed complexes was assessed through 100ns Molecular Dynamics (MD) simulations using Gromacs-2022.2 with the AMBER99SB-ILDN [17] force field. Since MD simulations are both time- and resource-intensive for most complexes, we opted to evaluate only the best-designed candidate, i.e., the one with the lowest RMSD when structurally compared to its native counterpart out of the pool of eligible candidates. In the MD simulation process, the complex structure is first solvated in a cubic box with atoms 1.0nm from the edges, and the solvated systems are made electrically neutral by adding the necessary number of positive or negative ions. Next, energy minimization of the system is done using 1000 steps of the steepest descent algorithm. The system is then equilibrated for 100ps at 298 K to stabilize temperature, then for 1 ns at 298 K to reach 1 bar pressure. Finally, a 100ns simulation is run at 298 K.

### 2.3 Existing comparable methods set-up

While comparing with five existing methods, (viz. data-driven Deep Learning-based protein design method (TIMED [4]), enerative model-based approach (GENERALIST [1]), evolution-based method (EBM [23]), and approaches using MCM (Modularity-based [24] and SADIE [20])), we provide either the sequence or structure of the native mutating chain (from the chain-pair of a protein complex) as input and obtain the designed candidates, which were then passed through the designed sequences’ verification pipeline in our method, up to complex structure prediction by AlphaFold2. The designed complex structures obtained by AlphaFold2 prediction were then compared with the native complex structure to find the RMSD. We also noted the pTM [14] and ipTM [8] values as given by AlphaFold2 as a part of complex structure prediction. For TIMED, we input the native mutating chain structure in PDB format to its pre-trained model for the inference and finally yielded a number of generated sequences through sampling. In the case of SADIE, while it originally started with random decoy sequences, we used the native mutating chain sequence. For GENERALIST, the native mutating chain sequence in FASTA format was used to generate multiple designed sequences through inference.

### 2.4 PU learning

#### Dataset preparation

The positive data points for this approach were sourced from the native protein complexes of the Protein-Protein Docking Benchmark and the results of the MD simulation. Negative data points were derived from the MD simulation results as well as from other steps, such as chain structure and complex structure comparisons. Unlabeled data points came from the PMCM simulation’s accepted data points, which were not part of the HDBSCAN clustering results. We structured the PUL dataset so that approximately 90.5% of the total data points are unlabeled, around 1.5% are positive, and roughly 8% are negatively annotated, proportionately higher than that of positive candidates. In selecting the unlabeled data points accepted from the PMCM simulation, we adhered to the specified percentage composition. For each data point, represented by a pair of sequences corresponding to the fixed and designed chains, we utilized PLM-based embeddings, such as those from ProtTrans [5] (ProtT5). These embeddings were generated for each sequence and then subjected to Principal Component Analysis (PCA) for dimensionality reduction. The PLM embeddings for each chain sequence were previously calculated during the PMCM simulation process for the MaTPIP-based [10] prediction, so we reused them instead of recalculating them. Additionally, we incorporated the PPI score and the number of mutations in the designed chain as extra features for each data point. For the native protein complexes sourced directly from the Docking Benchmark, the number of mutations was set to zero (indicating no mutations).

#### Selection of KAN as classifier

We applied sample-selection-based PU learning [2], which involves: (i) analysis of reliable negative samples as identified using a classifier; (ii) training of a supervised classifier on a dataset containing both positive and reliable negative samples. We had opted for a KAN-based classifier [18, 11] over MLP for better accuracy, smaller model, and better network expressiveness. KAN can converge faster than MLP if there exists a KA (Kolmogorov-Arnold) representation of the underlying function (as shown in the KAN paper [18]).

#### Training and evaluation

For training and evaluating the PU learning model, we implemented a custom 3-fold cross-validation (CV). First, we split the entire dataset into two groups: one containing only positive and negative samples and another with only unlabeled samples. The positive and negative sample dataset was then divided into 3 stratified folds, ensuring each fold maintained the class ratio. The *i*^*th*^ fold (*i* = 0, 1, 2) was treated as the test set, while the other two were combined and further underwent a stratified split into a 9:1 ratio to form training and validation sets. In the training set, all negative samples were removed and replaced with the unlabeled samples (from the dataset containing only unlabeled samples), ensuring training occurred only on positive and unlabeled samples, while both validation and test sets contained positive and negative samples. To address class imbalance during training, we applied class-weight techniques, assigning higher weights to the positive class in the loss function to penalize misclassifications more heavily.

All experiments were conducted on a machine with 187 GB RAM, a 16 GB Nvidia Tesla V100 GPU, and an Intel Xeon Gold 6148 CPU @ 2.40 GHz. The training dataset size was nearly 2 GB.

### 2.5 Implementation details

Configuration for KAN-classifier training includes the following: Initial learning rate = 0.001 (used with Adam optimizer); ExponentialLR’ learning-rate scheduler was used with gamma = 0.8; Number of epochs = 20; grid size = 5; spline order = 3; scale spline = 1.0; scale noise=0.1. For AlphaFold, we utilized a local implementation of ColabFold [22] referred to as LocalColabFold. We executed MD simulation using Gromacs-2022.2 with the AMBER99SB-ILDN [17] force field.

## 3 Results and Discussion

### 3.1 Sequence synthesis results of PMCM

For a protein complex with a mutating chain length of approximately 300, the average simulation time per iteration (PMCM) is under 1 second (and would likely be even shorter in a dedicated runtime environment). This indicates that both types of simulations are highly efficient in generating the designed chain sequences. It is observed that around 20% of the iterations were accepted for almost all protein complexes. Too high (*>* 90%) or too low (*<* 5%) acceptance rate would require further investigation of the setup or the effectiveness of the proposed simulation approach. Another key observation is that, for most complexes, the PPI score (accepted only if above 0.5) fluctuated within the probability range [0.5, 1.0] rather than remaining confined to a specific section of this range. This suggests that the perturbations introduced by the Metropolis criterion were successful to some extent in preventing convergence to local optima. It was anticipated that for each protein complex in PMCM, certain intervals within the [0.5, 1.0] range would show higher accumulations, but the Metropolis criterion would ensure entries across other intervals, though proportionally fewer. The majority of complexes met this expectation, as demonstrated by the PMCM results. For instance, some cases exhibited a significant accumulation in the upper interval [0.9, 1.0], while smaller entries were also present in other intervals. A similar pattern was observed in the lower interval [0.5, 0.6] for certain cases.

### 3.2 Verification (AlphaFold2 and MD simulation) results

AlphaFold2 predicted structures of the designed protein complexes are compared with native complexes, resulting in a median RMSD of 1.17 Å. The median pTM [14] and ipTM [8] scores, used as confidence metrics from AlphaFold2 for the designed complex structure predictions, are 0.72 and 0.88, respectively.

#### Result comparison

Five different methods were compared with our proposed approach. In Fig 2A, the RMSD values (between the designed and native protein complex structures) are presented in box plots for each method. Fig 2B, and 2C show the method-wise box plots of pTM and ipTM scores, respectively. In all plots, we focused on the best-designed candidate protein complex, defined as the one with the lowest RMSD value when compared to the native structure. The following abbreviations were used for result comparison: Our Method (PM), TIMED (TD), GENERALIST (GL), EBM-Design (ED), SADIE (SD), and Modularity-based (MB). For RMSD, lower values indicate better performance, while for pTM and ipTM, higher values are preferred. The pTM score reflects how accurately AlphaFold-Multimer [8] predicts the overall structure of a complex, whereas the ipTM score evaluates the accuracy of subunit positioning. A pTM score exceeding 0.5 and ipTM score above 0.8 are preferred.

**Fig. 2.**
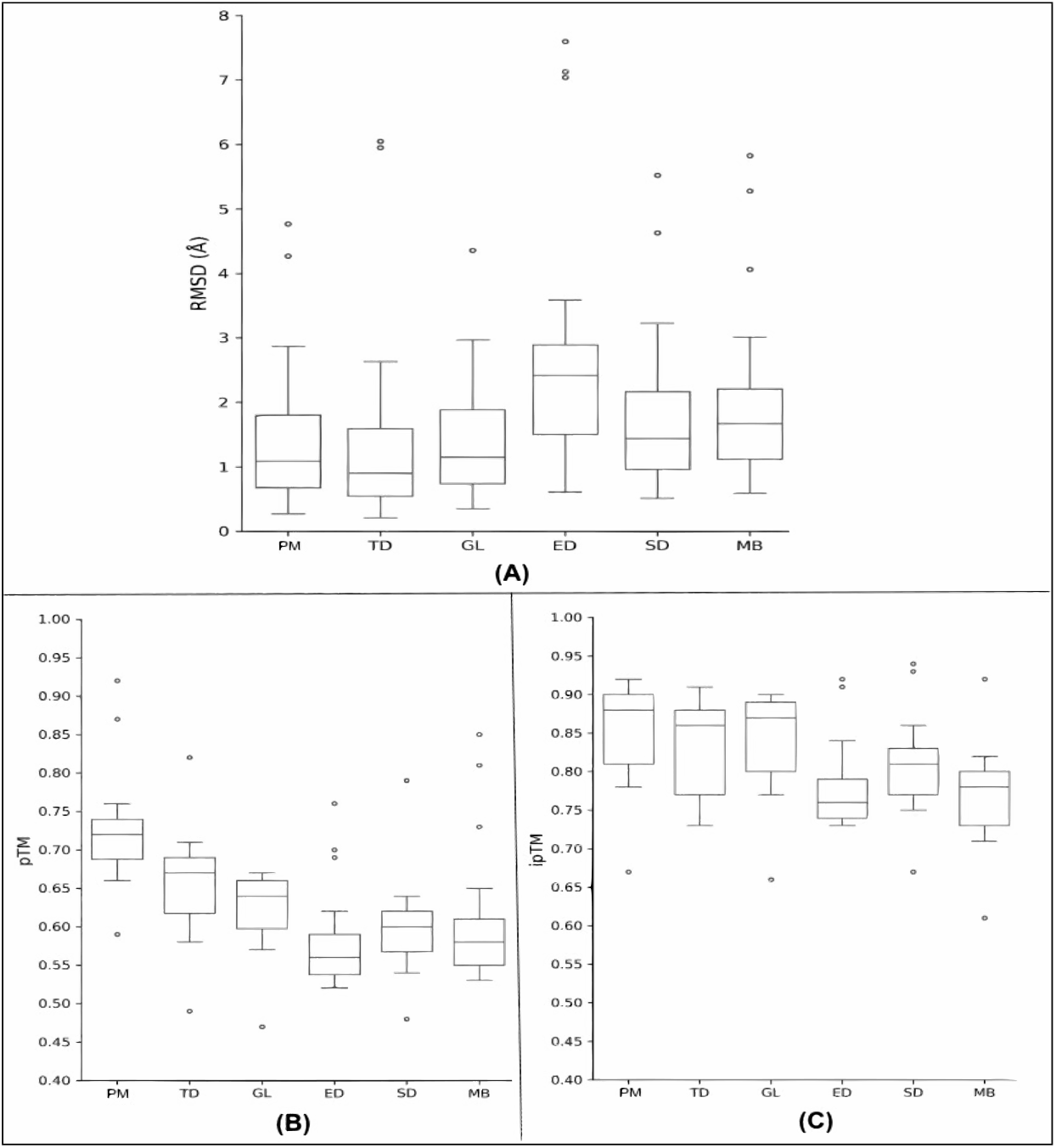
Box plot comparisons with other methods. RMSD values between predicted designed and native protein complex structures, pTM values and (C) ipTM values, given by AlphaFold2 as confidence metrics for designed protein complex structure predictions. For RMSD, the lower the value is better, but for pTM and ipTM, the higher is better. RMSD closer to zero, pTM ≥ 0.5, and ipTM ≥ 0.8 are preferred. **Abbreviations used for the name of the methods along the X-axis:** Our Method (PM); TIMED (TD); GENERALIST (GL); EBM-Design (ED); SADIE (SD); Modularity-based (MB);

From Fig 2A, 2B, and 2C, it is evident that our proposed method (PM) shows a slightly better median performance across all three metrics (RMSD, pTM, and ipTM) compared to GL. Additionally, PM outperforms ED, SD, and MB by a significant margin in all cases. While the median RMSD of TD is slightly better than that of PM (Fig 2A), PM has a higher median pTM and ipTM scores (Fig 2B, 2C), indicating that although TD has a marginal edge in RMSD, PM provides higher prediction confidence according to AlphaFold2.

Another important observation is that the box plots for PM, TD, and GL in Fig 2A are positively skewed, indicating that more RMSD values are concentrated in the lower range, which is desirable. The box plot for SD shows slight positive skewness, while the box plots for MB and ED exhibit different patterns—MB is symmetrical, and ED is negatively skewed, implying more RMSD values concentrated in the higher range (undesirable). For pTM and ipTM (Fig 2B and 2C), a reverse trend is observed: PM, TD, and GL show desirable negative skewness with higher concentrations of pTM values in the upper range. SD shows slight negative skewness, MB remains symmetrical, and ED shows an unfavorable positive skewness with more pTM values in the lower range. For ipTM (Fig 2C), all methods except ED display the desired negative skewness.

In summary, this comparative analysis demonstrates that the our proposed approach clearly excelled over the standard evolutionary methods (EBM) and recent simulation-based approaches (SADIE, Modularity-based) while remaining highly contending with the latest data-driven, Deep Learning-based protein design techniques (TIMED, GENERALIST).

#### AlphaFold2 metrics

The predicted local distance difference test (pLDDT) [14] provides a per-residue measure of local confidence, ranging from 0 to 100, with higher values indicating greater confidence and typically more accurate predictions. Predicted aligned error (PAE) [8] assesses how confident AlphaFold2 is in the relative positioning of two residues within a predicted structure. A low PAE score between residues from different domains suggests low predicted error, meaning AlphaFold2 is confident in their positions. Since both pLDDT and PAE are calculated at the residue level for each complex, any aggregated presentation of these metrics at the complex level for a group of designed complexes would be less meaningful.

#### MD simulation result

Given that the 100 ns MD simulations are highly time-consuming, we focused primarily on assessing the structural stability of the best-designed complexes (chosen from a pool of at least four eligible candidates per native complex). Even with this limited scope, we found that in nearly 73% of the cases, the average RMSD remained below 2 *Å*, indicating structural stability. This percentage could likely be improved on the second or third-best candidates for those complexes where the top candidate exceeded 2 *Å* in average RMSD. Fig 3 illustrates RMSD, RMSF, and Rg plots (designed versus native complexes) derived from the 100 ns MD simulations for a few selected cases.

**Fig. 3.**
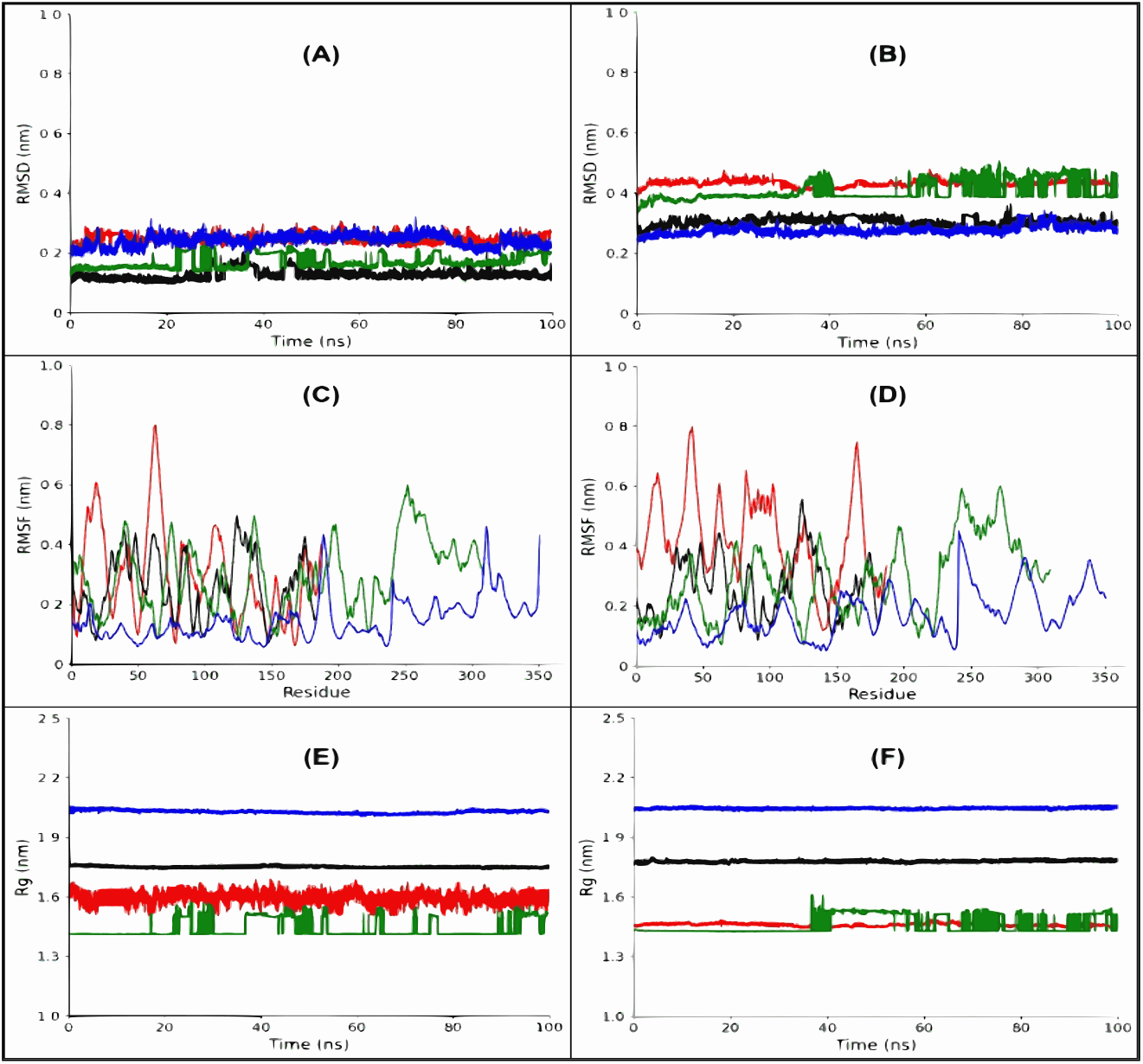
RMSD (A: native, B: design), RMSF (C: native, D: design), Rg (E: native, F: design) plots from 100ns MD simulations for a few protein complexes. Color code for the protein complex with PDB ID (1KTZ, red); (1AY7, black); (1UDI, green); (1YVB, blue). The results show that for RMSF and Rg, the designed candidates are closely following their native counterparts, whereas for RMSD, there are slight gaps for a few cases, but the designed candidates’ RMSDs remain well below 2*Å*.

#### Interface residue mutation for altered interaction

We did not limit mutations to only interface residues to change the interaction; instead, we applied mutations across the full chain sequence to ensure the stability of the newly designed protein complex. In all the selected designed sequences, interface residues were also mutated.

### 3.3 PU learning result

The KAN-based classifier notably outperformed RFC in all metrics and MLP-based classifier, except recall (Table 1). In our PU learning application, precision is more critical than recall because false positives are costlier than false negatives. A false positive could lead to wasted resources, as it would trigger costly experimental verification, which would eventually fail. However, while reducing false negatives is important, it does not directly result in wasted resources. Thus, higher precision (fewer false positives) is more desirable than higher recall (fewer false negatives). Table 1 also suggests that the KAN-based PU learning classifier can effectively replace significant portions of the resource-intensive designed candidate selection and verification stages in our method (e.g., HDBSCAN-based clustering and AlphaFold2-based structure prediction). To further enhance the performance of the KAN-based classifier and potentially replace even the MD simulation process, more labeled data, is required for the training set. However, the performance of the KAN-based PU learning classifier is enough for the selection and verification pipeline to replacing more resource-consuming processes.

**Table 1.**
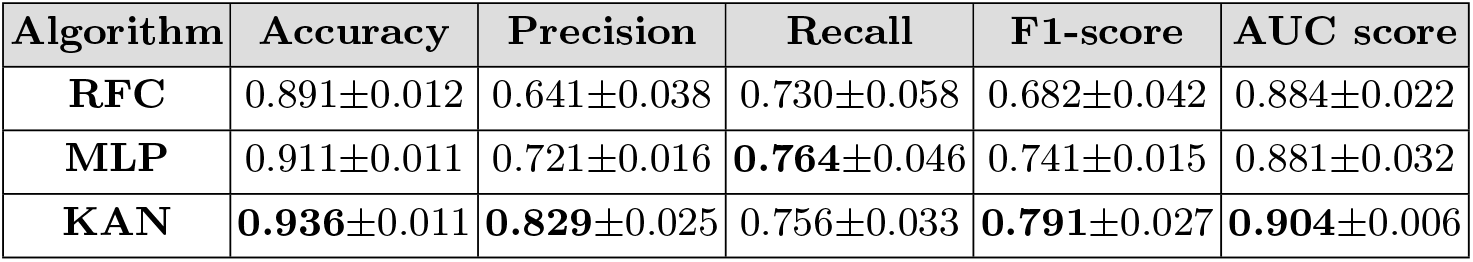
PU learning result. Each performance metric in the table is reported as mean±standard deviation across the custom 3-fold CV. **Bold** indicates best values.

## 4 Conclusion

In summary, we introduce a range of novel concepts in protein design that can enrich the overall protein interaction engineering process. Our PMCM uses deep learning model MaTPIP-Hybrid [10] for sequence generation. The use of PU learning replaces the resource-heavy designed sequence selection and verification pipeline, and the KAN-based approach for PU learning, had demonstrated consistent, and often superior performance compared to other methods, including Deep Learning-based methods. The proposed simulation-based methods (PMCM) are relatively more non-deterministic or randomized compared to learning-based approaches. However, our proposed protein interaction design approach convincingly outclassed traditional evolutionary and recent simulation-driven methods and remains competitive with the latest data-driven, Deep Learning-based protein design techniques. Future work may evaluate advanced PLMs such as ESM-2 [16] and further enhance the KAN-based PU classifier by adopting alternative PU learning techniques or incorporating additional labeled data into the training set. Overall, this work presents a range of innovative ideas in the field of protein interaction design, laying the groundwork for future research endeavors.

## Acknowledgments

The work is partially supported by the IIT Kharagpur AI4ICPS I Hub Foundation Fund, IIT Kharagpur Campus, West Bengal - 721302, India (Sanction Letter No: TRP3RD70001, Dt. 20-02-2024) and the Department of Science and Technology and Biotechnology, Government of West Bengal, Vigyan Chetana Bhavan, Salt Lake, DD 26/B, Sector-I, Kolkata - 700064 (Sanction Letter No: 2053(Sanc.)/STBT-11012(31)/1/2024-ST SEC, Dt. 31-01-2024).

